# Low-dose lung radiotherapy for COVID-19 lung disease: a preclinical efficacy study in a bleomycin model of pneumonitis

**DOI:** 10.1101/2021.03.03.433704

**Authors:** Mark R Jackson, Katrina Stevenson, Sandeep K Chahal, Emer Curley, George E Finney, Rodrigo Gutierrez-Quintana, Evarest Onwubiko, Angelika F Rupp, Karen Strathdee, Megan KL MacLeod, Charles McSharry, Anthony J Chalmers

## Abstract

**Purpose:** Low-dose whole lung radiotherapy (LDLR) has been proposed as a treatment for patients with acute respiratory distress syndrome associated with SARS-CoV-2 infection and clinical trials are underway. There is an urgent need for preclinical evidence to justify this approach and inform dose, scheduling and mechanisms of action.

**Materials and methods:** Female C57BL/6 mice were treated with intranasal bleomycin sulphate (7.5 or 11.25 units/kg, day 0), then exposed to whole lung radiation therapy (0.5, 1.0, 1.5 Gy or sham, day 3). Bodyweight was measured daily and lung tissue harvested for histology and flow cytometry on day 10. Computed tomography (CT) lung imaging was performed pre-radiation (day 3) and pre-endpoint (day 10).

**Results:** Bleomycin caused pneumonitis of variable severity which correlated with weight loss. LDLR at 1.0 Gy was associated with a significant increase in the proportion of mice recovering to 98% of initial bodyweight and a proportion of these mice exhibited less severe histopathological lung changes. Mice experiencing moderate initial weight loss were more likely to respond to LDLR than those experiencing severe initial weight loss. Additionally, LDLR (1.0 Gy) significantly reduced bleomycin-induced increases in interstitial macrophages, CD103+ dendritic cells and neutrophil-DC hybrids. Overall,bleomycin-treated mice exhibited significantly higher percentages of non-aerated lung in left than right lungs and LDLR (1.0 Gy) prevented further reductions in aerated lung volume in right but not left lungs. LDLR at 0.5 and 1.5 Gy did not modulate bodyweight or flow cytometric readouts of bleomycin-induced pneumonitis.

**Conclusions:** Our data support the concept that LDLR can ameliorate acute inflammatory lung injury, identify 1.0 Gy as the most effective dose and provide preliminary evidence that it is more effective in the context of moderate than severe pneumonitis. Mechanistically, LDLR at 1.0 Gy significantly suppressed bleomycin-induced accumulation of pulmonary interstitial macrophages, CD103+ dendritic cells and neutrophil-DC hybrids.

## Introduction

To date (February 2021), infection with the Severe Acute Respiratory Syndrome Coronavirus 2 (SARS-CoV-2) has been associated with more than 2,500,000 deaths globally (1). Infection by SARS-CoV-2 can cause a clinical syndrome termed COVID-19 which varies widely in severity with a small proportion of patients developing severe pneumonia and life-threatening acute respiratory distress syndrome (ARDS) (2). COVID-19 lung disease is characterised by pathological inflammation, the severity of which correlates with morbidity and mortality (3). The characteristic lung pathology and associated systemic deterioration are consistent with ARDS and cytokine release syndrome (CRS) respectively (4). Key features include florid alveolar infiltration by neutrophils, macrophages and lymphocytes and markedly increased pro-inflammatory cytokines including interleukin-6 (IL-6), IL-1β, tumour necrosis factor (TNF) and interferon gamma (IFNγ) (5,6).

Treatment options for COVID-19 remain limited. The open-label RECOVERY trial (UK) reported that dexamethasone reduced 28-day mortality compared with standard of care among hospitalised patients who required ventilatory support (Relative risk [RR] 0.65) or oxygenation (RR 0.80) (7). More recently, the REMAP-CAP trial reported improvements in survival and time to recovery following dual therapy with tocilizumab and sarilumab (unpublished data), and the anti-inflammatory agent colchicine has been reported to reduce hospitalisation and mortality in patients with COVID-19 infection who had a least one risk factor for complications (8). While some of these studies await peer review, the early efficacy data support the concept of acute inflammation being the critical pathological process in COVID-19 lung disease, and indicate that broad spectrum immunosuppressive therapies may be of therapeutic value.

Low-dose whole lung radiotherapy (LDLR; radiation doses of 1.5 Gy or less) was used extensively as a treatment for pneumonias of various aetiologies in the pre-antibiotic era (9). In various preclinical models of inflammation, LDLR has been shown to induce anti-inflammatory cytokine production, reduce leucocyte-endothelial adhesion, and repolarise myeloid and lymphoid cells towards immune-suppressive phenotypes (10). Recent preclinical studies have generated preliminary data to indicate that LDLR (0.5-1 Gy) reduces pneumonitis in lipopolysaccharide (LPS) and influenza virus mouse models (11). These diverse but limited bodies of evidence have underpinned early phase clinical trials of LDLR for acutely unwell COVID-19 patients in several countries including the USA, India and Spain. The safety of the intervention is supported by preliminary results from phase I trials (12,13), in which early signals of efficacy were reported including in very elderly comorbid patients with severe COVID-19 (12).

This therapeutic approach has generated intense controversy (14-17). While expressing diverse opinions, the vast majority of stakeholders have emphasised the urgent need for high quality preclinical data to (i) justify (or not) the commencement of clinical studies, (ii) elucidate mechanisms of efficacy and (iii) inform decisions on radiation dose, scheduling and target volume (18). To address this need, we undertook preclinical studies using a mouse model of bleomycin-induced pneumonitis. Many pathophysiological changes of COVID-19 lung disease (epithelial cytopathy, endotheliitis, inflammatory infiltrates, surfactant loss) are reproduced in the pneumonitis induced by inhaled bleomycin (19,20). Indeed, single cell sequencing studies of mouse bleomycin (21) and COVID-19 (22,23) pneumonitis have shown pathogenic SPP1^pos^macrophage orthologues expressing key inflammatory mediators to be prominent in both conditions, along with profoundly reduced expression of anticoagulant and anti-apoptotic protein S in alveolar macrophages. Exogenous protein S is protective of bleomycin pneumonitis (24) and has been proposed as a potential treatment for COVID-19 (25).

To test the hypothesis that LDLR would reduce the severity of bleomycin-induced acute lung injury by exerting suppressive effects on cellular and molecular components of the inflammatory response, we measured the effects of 0.5, 1.0 and 1.5 Gy whole lung irradiation on bodyweight (primary endpoint); pulmonary cytology and histology, and lung computed tomography (CT) appearances (secondary endpoints). Our data show that 1.0 Gy LDLR enhances recovery in a proportion of bleomycin-treated mice, with corresponding improvements in lung histopathology and imaging parameters and modulation of specific immune cell populations.

## Materials and Methods

### Reagents

All reagents were purchased from Biolegend unless otherwise stated. Bleomycin sulphate was obtained from European Pharmacopeia EDQM, Council of Europe, France. Intranasal bleomycin dose was 11.25 units/kg except in the initial pilot study, when 7.5 units/kg was also used.

### Experimental pneumonitis

Bleomycin generates a well-established murine model of pneumonitis (20) with a dynamic pathology similar to that of COVID-19. Female, 11-13 week old C57BL/6 mice (Charles River Laboratories) were administered one intranasal 40 μL dose of bleomycin sulphate (7.5 or 11.25 units/kg) or phosphate buffered saline (PBS) vehicle control under light isoflurane anaesthesia. Mice were maintained in a pathogen-free facility, provided with additional high calorie, soft diet to ease feeding, and monitored daily for wellbeing and change in bodyweight. Those demonstrating signs of illness such as lethargy, isolation, reduced mobility, altered respiration, or ≥25% weight-loss were humanely culled. The experimental design optimised mice numbers to comply with the principles of Replacement, Reduction and Refinement for humane animal research. Procedures were governed by the Animals Scientific Procedures Act 1986 and approved by Home Office licence PP6245051.

### Low-dose lung radiotherapy (LDLR)

Bleomycin-treated mice exhibiting a day 3 relative bodyweight area-under-the-curve (AUC) ≤2.92 were randomised to receive LDLR or sham irradiation. Anaesthetised mice were irradiated with low-dose whole lung radiotherapy (0.5, 1.0, 1.5 Gray or sham) on day 3, using the Small Animal Radiation Research Platform (SARRP) developed by XStrahl. A 220 kVp, 13 mA X-ray beam was used with a dose rate of approximately 280 cGymin^-1^ at the chosen aperture size. Treatment was delivered with anterior and posterior parallel opposed fields. The broad focal spot (5.5 mm) was used and the SARRP’s motorised variable collimator set to an aperture size of 20 x 20 mm to ensure full coverage of both lungs. Mice were culled on day 10 and lung tissue harvested for experimental endpoint analysis as described below.

### CT assessment of lung changes

Lung changes were measured using the SARRP’s in-built cone-beam CT (CBCT) function to image anaesthetised mice on days 3 (pre-irradiation) and 10 (experimental endpoint). Images were reconstructed using the FDK (Feldkamp, Davis and Kress) CBCT reconstruction algorithm from 1440 projections taken at 60 kVp and 0.8 mA using the fine focal spot (1 mm). For quantification of aerated lung volumes, Hounsfield unit (HU) clinical ranges were used: poorly aerated lung defined as -500 to -100 HU and normo-aerated defined as -900 to -500 HU. Images were analysed using the Lung CT analyzer module from the 3D Slicer software extension SlicerCIP (26,27).

### Tissue collection

Mice were culled by terminal intraperitoneal injection of 100 μL sodium pentobarbital (200 mg/ml) and cardiac exsanguination. The trachea was exposed, a small transverse opening cut between cartilage rings and a ligature tied loosely distal to the cut. The protruding 0.5 cm tip of a cannula sheath around a 23G syringe needle was inserted into the opening and the ligature tightened. The lungs were lavaged twice with 0.8 mL PBS and then perfused via the right ventricle with cold PBS until they blanched, after which lungs and heart were removed *en bloc*. The left lobe of the lung was excised, submerged in 4% neutral buffered formalin fixative for 24hr and processed for histology while the right lung lobes were processed for cytology.

### Lung histology

Serial 4 μm sections of the left lobe were cut and stained with haematoxylin and eosin (H&E) and Masson’s trichrome and evaluated independently by a veterinary pathologist and a pulmonary immunologist both of whom were blinded to the experimental treatment. In brief, semiquantitative scoring (described in detail in Supplementary data) examined the extent of interstitial mononuclear cell infiltrates, specifically interstitial (to intra-alveolar) macrophage infiltrates and perivascular/peribronchiolar lymphocyte aggregates.

### Lung tissue cytology

Small pieces of right lung were incubated with dispase (3.2 mg/mL, Roche), collagenase P (0.4 mg/mL, Roche), and DNAse I (0.2 mg/mL, Sigma) in 2 mL RPMI at 37°C in a shaking incubator for 40 minutes. Lung pieces were transferred into a 100 mm strainer and a single cell suspension prepared and filter rinsed to transfer all cells into a 50 mL tube. Red blood cells were lysed by RBC lysis buffer (ThermoFisher) and viable cells counted. Next, cells were Fc blocked with 24G2 antibody and normal mouse serum for 10 minutes then incubated with fluorescently labelled antibodies for 20 minutes at 4°C. Antibodies are listed in Supp. Table S1. Following PBS washing, cells were incubated with viability dye (Zombie Aqua) for 20 minutes at 4°C, washed with FACS buffer (PBS with 2% FCS, 2 mM EDTA and 5 mM sodium azide), fixed with 2% paraformaldehyde for 20 minutes at 4°C and washed again with FACS buffer then stored at 4°C. Flow cytometry data were acquired on a BD Fortessa and analysed using FlowJo (version10, BD Biosciences). The gating strategy (28) is shown in Supp. Fig. S1.

### Statistical analysis

Statistical analyses were performed in R 3.6.3 (29) using the packages “MESS” (30), “survival” (31) and “survminer” (32). Box and whiskers were plotted according to the Tukey method. Statistical tests used and group sizes are indicated for each experiment. Additional detail on use of area-under-the-curve (AUC), recovery probability and aerated lung volume analyses is presented as Supplementary information.

## Results

Initially, pilot studies were conducted to characterise the bleomycin-induced pneumonitis model and establish optimum dosing and scheduling parameters. Using mouse bodyweight as a marker of systemic response, we observed variable responses to intranasal bleomycin (Fig. 1A), as reported in other studies. Area-under-the-curve (AUC) analysis revealed that, despite visible effects at day 3 (Supp. Fig. S2), by day 10 the bodyweight of mice treated with 7.5 units/kg bleomycin was not significantly different to controls (Fig. 1B). In contrast, administration of 11.25 units/kg induced progressive weight loss in the majority of mice, with 25% exhibiting a severe reduction that triggered humane endpoint sacrifice but 25% failing to show a demonstrable response (Fig. 1A,B). Histological assessment on day 3 revealed multifocal, small, interstitial to intra-alveolar macrophage infiltrates and appreciable but small, perivascular and peribronchiolar lymphocyte aggregates in the majority of mice (Fig. 1C). These were accompanied by a robust reduction in alveolar macrophages and an increase in interstitial macrophages measured by flow cytometry in dispersed lung tissue (Fig. 1D).

**Figure 1:**
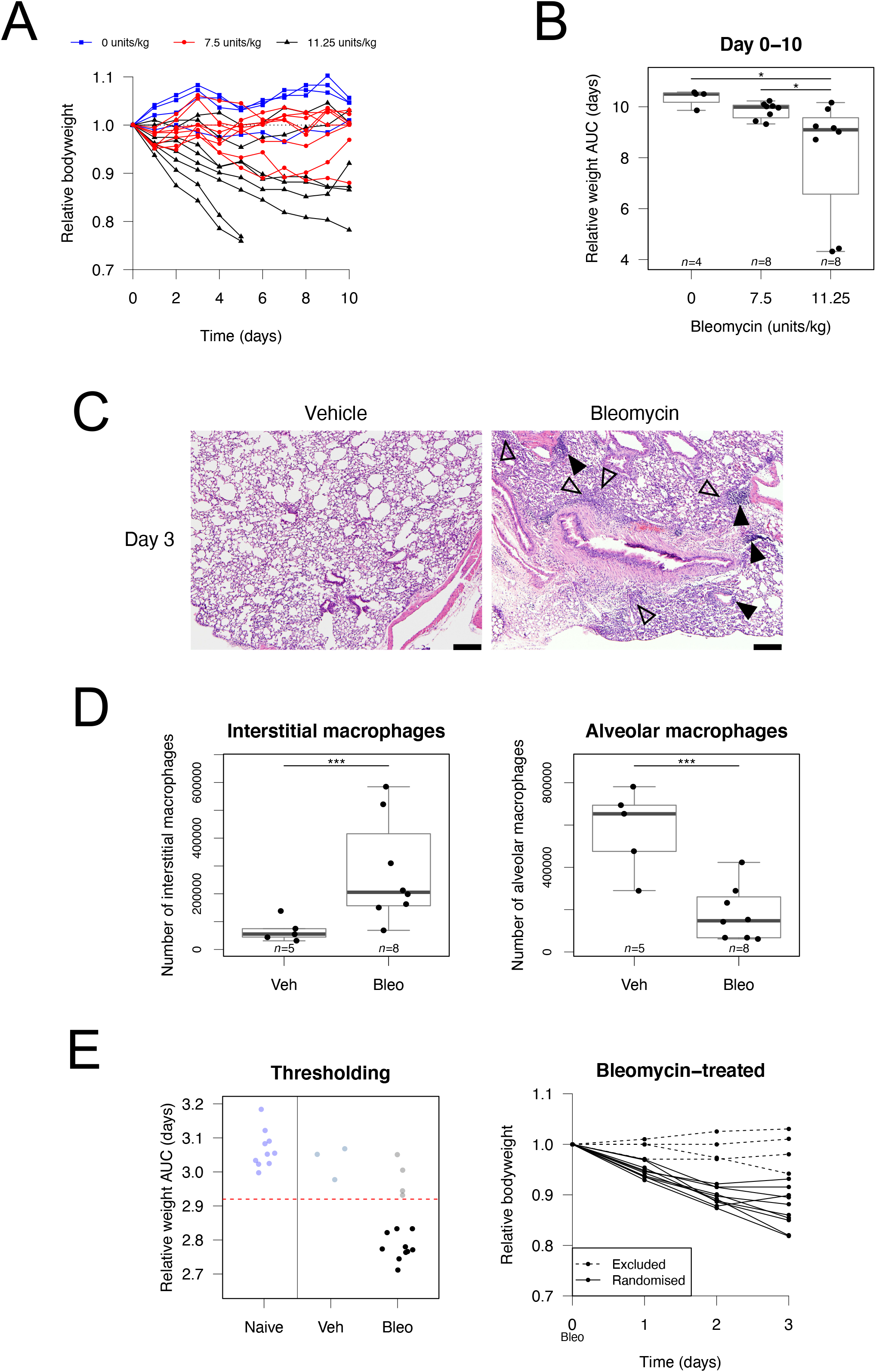
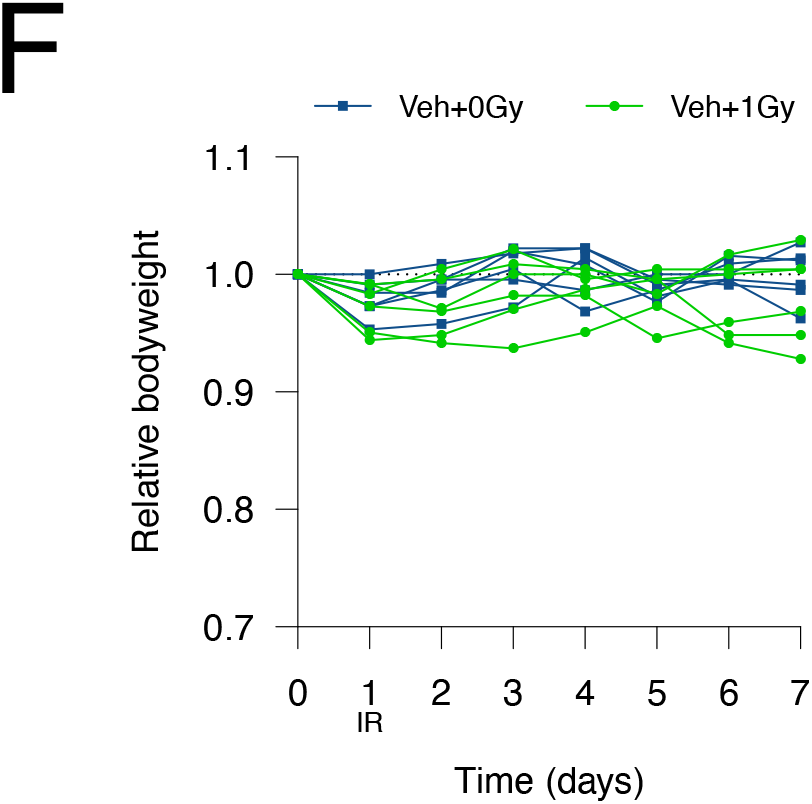
Bleomycin induces a variable pneumonitis in young adult, female C57BL/6 mice. (A) Relative mouse bodyweight following intranasal administration of bleomycin. (B) Area-under-the-curve (AUC) of relative mouse bodyweight up to day 10. (C) Pulmonary histology 3 days after treatment with vehicle (left) and bleomycin (11.25 units/kg, right). Macrophage infiltrates are annotated with open arrows and lymphocyte aggregates with filled arrows. Haematoxylin and eosin stain, scale bars: 200 μm. (D) Flow cytometric analysis of macrophages in mouse lung at day 3 following bleomycin treatment (11.25 units/kg). (E) Mice with day 3 relative bodyweight AUC ≤2.92 were selected for inclusion in efficacy studies. The corresponding relative bodyweights of excluded and included (randomised) mice are presented as an example. (F) Bodyweights of control mice treated with intranasal PBS then subjected to 1.0 Gy whole lung irradiation 24 hours later. Groups compared by Kruskal Wallis and *post hoc* Wilcoxon pairwise test, * P<0.05, *** P<0.001.

To mirror the clinical scenario, in which LDLR would only be considered in patients exhibiting moderate to severe COVID-19 lung disease, we opted to deliver LDLR three days after bleomycin treatment. Mice showing minimal weight loss at day 3 were excluded and those exhibiting a sustained drop in bodyweight (defined by day 3 AUC ≤2.92) were randomly allocated to receive LDLR or sham irradiation (Fig. 1E). Treatment of vehicle-only control mice with LDLR (1.0 Gy) was well tolerated with no effect on bodyweight and no detectable deviation from normal behaviour (Fig. 1F). Despite the variability inherent to the bleomycin model, treatment with 1.0 Gy was associated with a modest increase in mean bodyweight in irradiated versus sham- irradiated mice from day 5 onwards (Fig. 2A). Bodyweight plots for individual mice (Fig. 2B) illustrate the variable response to bleomycin and identify a sub-population of irradiated mice recovering to at least 96% of baseline bodyweight. Kaplan-Meier analysis demonstrated a statistically significant increase in the proportion of irradiated mice recovering to at least 98% of initial bodyweight after day 3 (21.2%, *n*=33) compared with sham-irradiated mice (3.3%, *n*=30; p=0.0265), with recovery also occurring earlier (Fig. 2C). This definition of recovery (regaining 98% of initial weight) was used as a reference in subsequent analyses. Recovery was also significantly increased in irradiated mice if a recovery threshold of 100% was imposed (p=0.0230) and a strong trend was observed at 96% (p=0.0776) (Table 1). Of note, treatment with LDLR did not increase the likelihood of an adverse outcome, with no difference in the proportions of irradiated and sham-irradiated mice experiencing severe weight loss (Fig. 2D, Supp. Fig. S3).

**Table 1:**
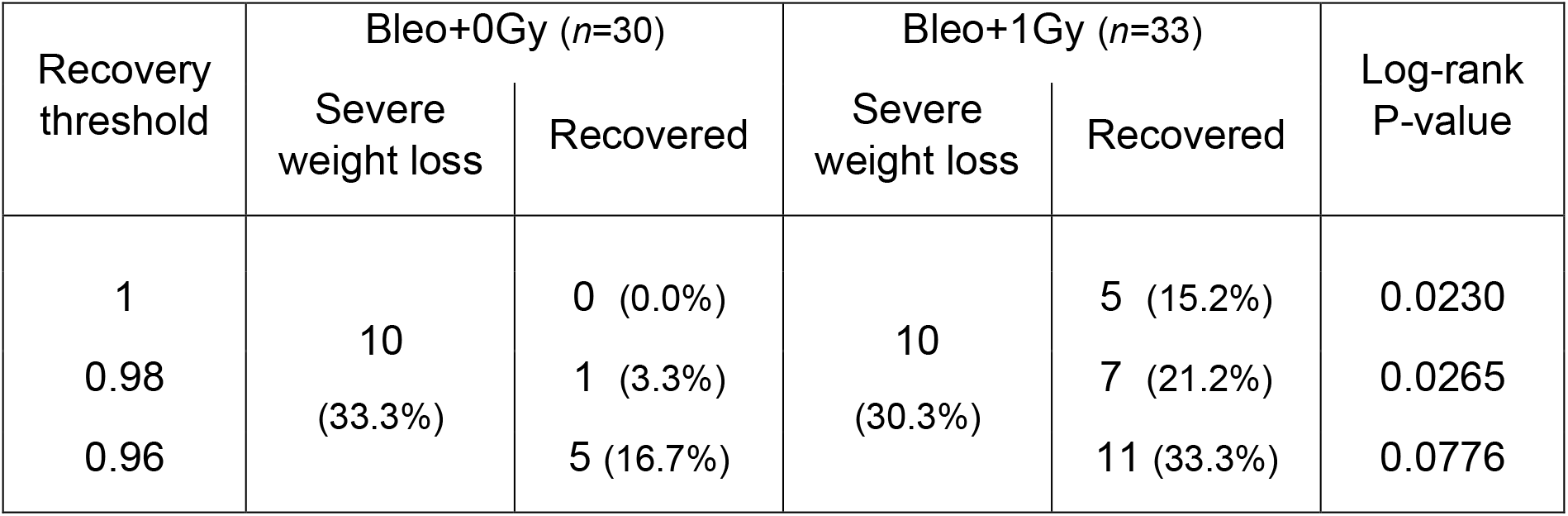
Log-rank analysis of recovery probability in mice treated with LDLR.

**Figure 2:**
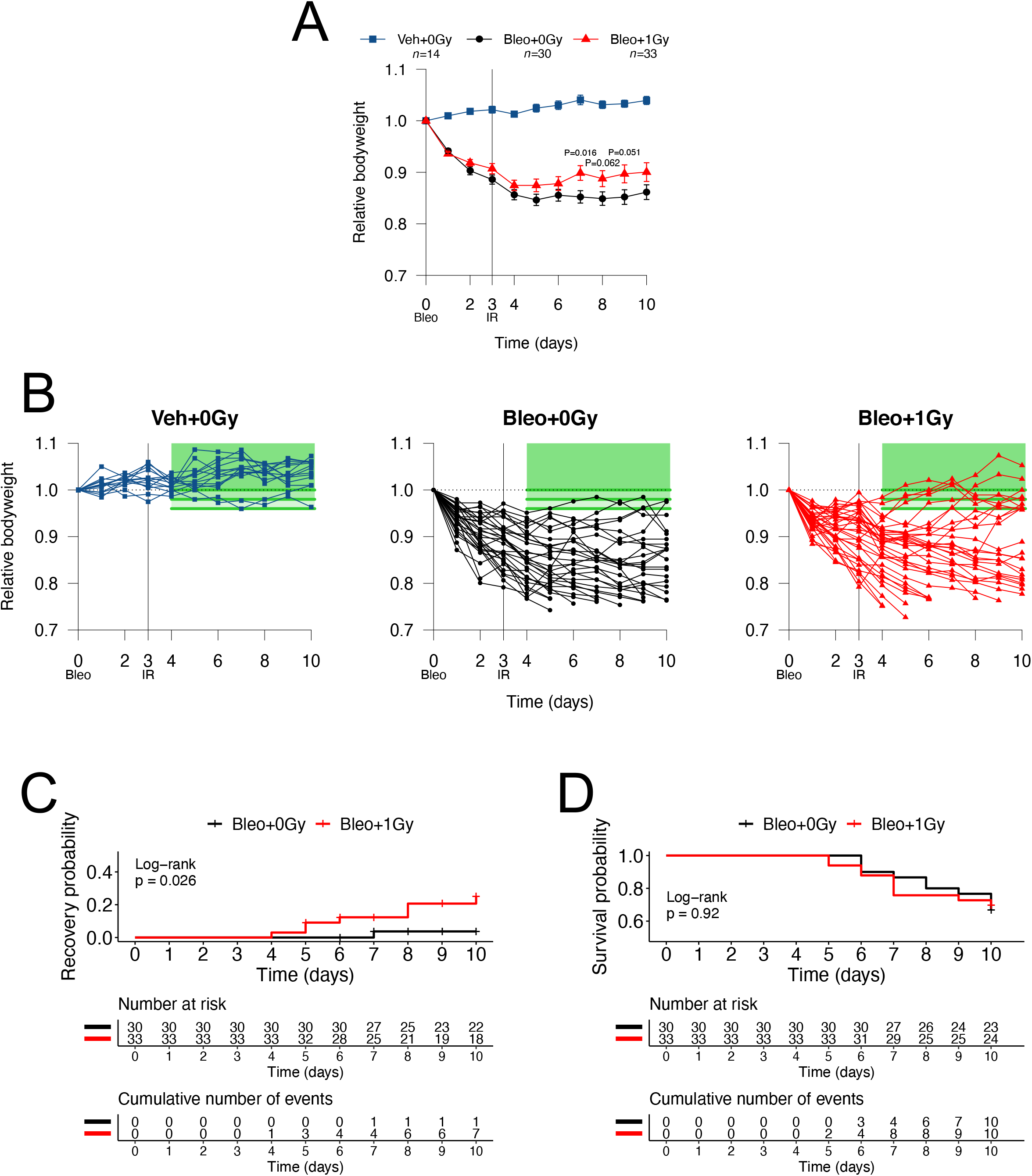
LDLR promotes recovery of bodyweight in a subset of bleomycin-treated mice. (A) Relative bodyweight (mean ± SEM) of bleomycin-treated mice treated with LDLR (1.0 Gy) or sham irradiation on day 3. Day-by-day comparison of sham and irradiated groups performed by t-test. (B) Relative bodyweight of individual mice; recovery defined as a return to 96, 98 or 100% of baseline bodyweight (green boxes) after day 3. (C) Kaplan-Meier analysis of recovery to 98% of baseline bodyweight; groups compared by log-rank test. (D) Kaplan-Meier analysis of mouse survival. Mice exhibiting severe weight loss were culled to comply with humane endpoint. Groups compared by log-rank test.

Histological assessment of mice on day 10 for macrophage infiltrates and lymphocyte aggregates demonstrated significant increases in immune infiltration of the lungs of bleomycin-treated mice compared to vehicle-treated mice (Fig. 3A,B). While no statistically significant difference in composite score was detected between irradiated and sham-irradiated groups, a subset of irradiated mice, composed predominantly of mice whose bodyweight had recovered after treatment (open symbols in Fig. 3A), exhibited lower levels of inflammatory cells. In keeping with this observation, there was a significant inverse correlation between histological composite score and relative bodyweight AUC across all mice exposed to bleomycin (*r*=-0.42, p=0.0048; Supp. Fig. S4). Early fibrotic changes were observed (Supp. Fig. S5) but were deemed not substantial enough to be quantified with existing scoring systems for fibrosis, which have been created and validated for later timepoints than those under investigation in this study.

**Figure 3:**
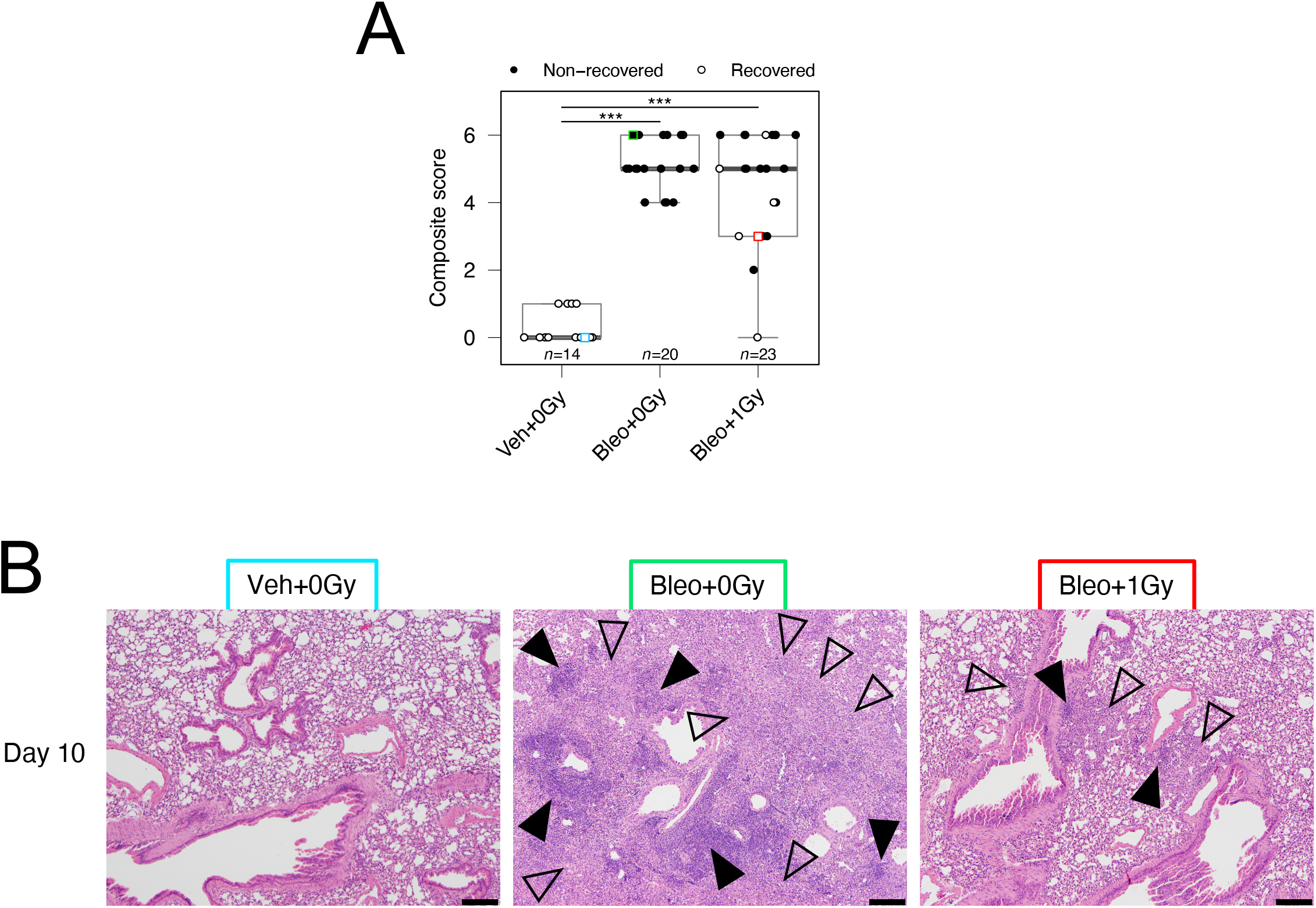
LDLR reduces severity of lung pathology in a subset of bleomycin-treated mice. (A) Histology composite scores of pulmonary macrophage infiltrates and lymphocyte aggregates at day 10. Square symbols indicate images presented in (B). Mice whose bodyweight returned to 98% of baseline were classified as recovered. Groups compared by Kruskal Wallis and *post hoc* Wilcoxon pairwise test, *** P<0.001. (B) Examples of pulmonary histology at day 10. Macrophage infiltrates are annotated with open arrows and lymphocyte aggregates with filled arrows. Images colour-coded for cross-referencing with square symbols in (A). Haematoxylin and eosin stain, scale bars: 200 μm.

Immunocytological assessment of mouse lungs on day 10 showed that while the bleomycin-induced increase in interstitial macrophages was significantly blunted by lung irradiation (Fig. 4A), the associated reduction in alveolar macrophages was not affected (Fig. 4B). Bleomycin associated increases in CD103+ dendritic cells and neutrophil-DC hybrids (33) were also significantly attenuated in mice exposed to 1.0 Gy LDLR (Fig. 4C,D). Representative FACS plots are shown in Supp. Fig. S6. In addition to changes in cell number, bleomycin inhalation was associated with increased expression of the co-stimulatory molecule CD86 on alveolar macrophages and on neutrophil-DC hybrids, but reduced expression on CD103+ dendritic cells and interstitial macrophages (Supp. Fig. S7). Importantly, the reduction in expression of CD86 induced by bleomycin in interstitial macrophages was significantly attenuated by LDLR.

**Figure 4:**
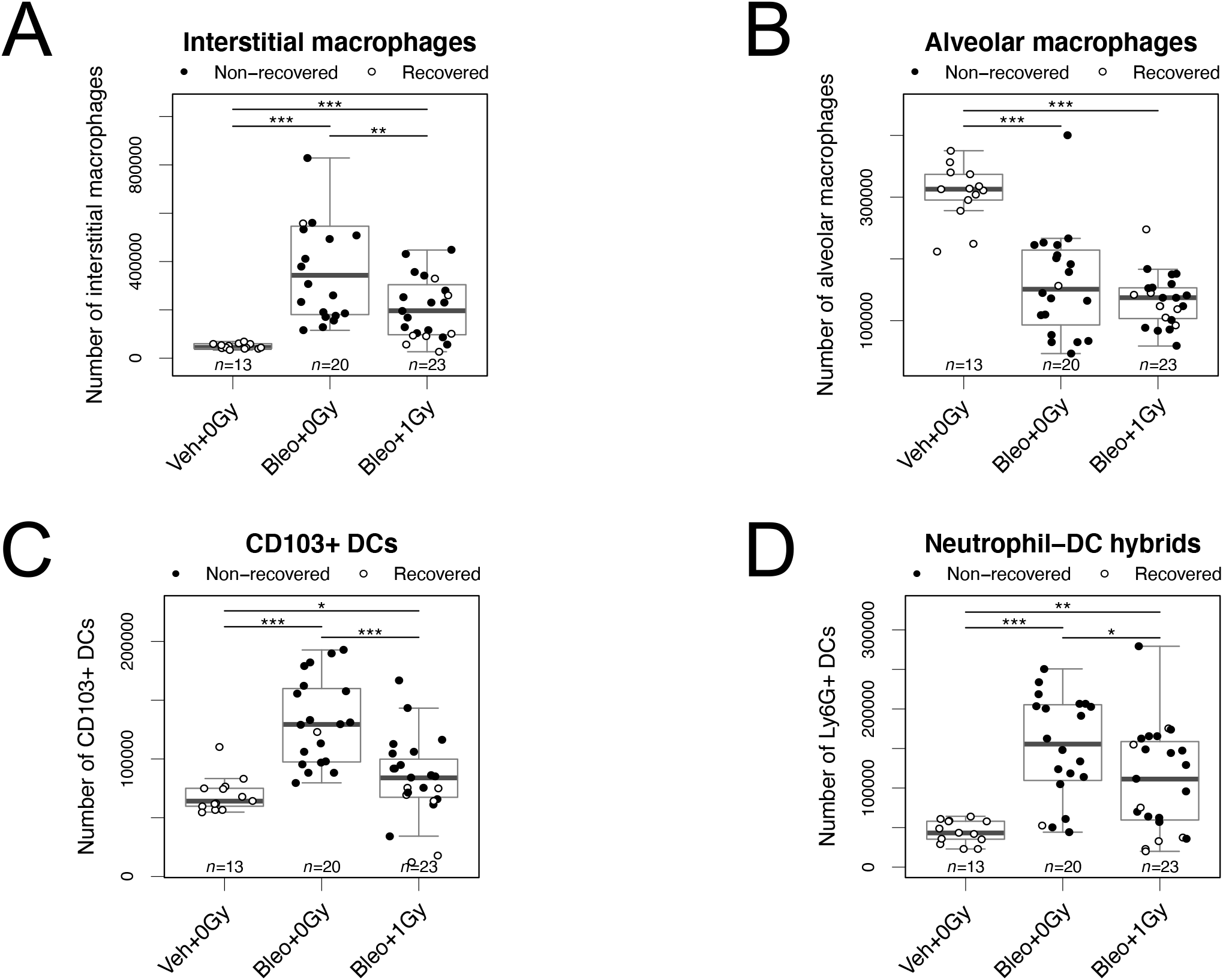
LDLR suppresses bleomycin-induced changes in immune cell numbers in mouse lung. Flow cytometric analysis of interstitial macrophages (A), alveolar macrophages (B), CD103+ dendritic cells (C) and Ly6G+ neutrophil-dendritic cell hybrids (D) in mouse lung at day 10 after bleomycin treatment. Mice whose bodyweight returned to 98% of baseline were classified as recovered. Groups compared by Kruskal Wallis and *post hoc* Wilcoxon pairwise test, * P<0.05, ** P<0.01, *** P<0.001.

To enable longitudinal assessment of lung infiltration, mice underwent CT imaging of the thorax on day 3 (pre-irradiation) and day 10. As previously reported, bleomycin related changes were significantly more pronounced in the left lung (Fig. 5A,B); this is thought to be due to morphological differences between left and right main bronchi (34). Left and right lung datasets were therefore analysed separately. Consistent with evolving acute lung injury, aerated lung volume decreased between days 3 and 10 in sham irradiated mice (both lungs) and in the left lungs of irradiated mice (Fig. 5C). In contrast, no statistically significant deterioration was observed in the right lungs of irradiated subjects (Fig. 5C, right panel). Furthermore the mean decrease in right lung aerated volume was significantly less in irradiated mice than controls (−3.8% and - 11.9% respectively, Fig. 5D). Indeed 36% (*n*=22) of irradiated mice showed an improvement (change >0%) in right lung aeration at day 10, compared to only 5% (*n*=19) of controls. No effect of irradiation was observed in the left lungs.

**Figure 5:**
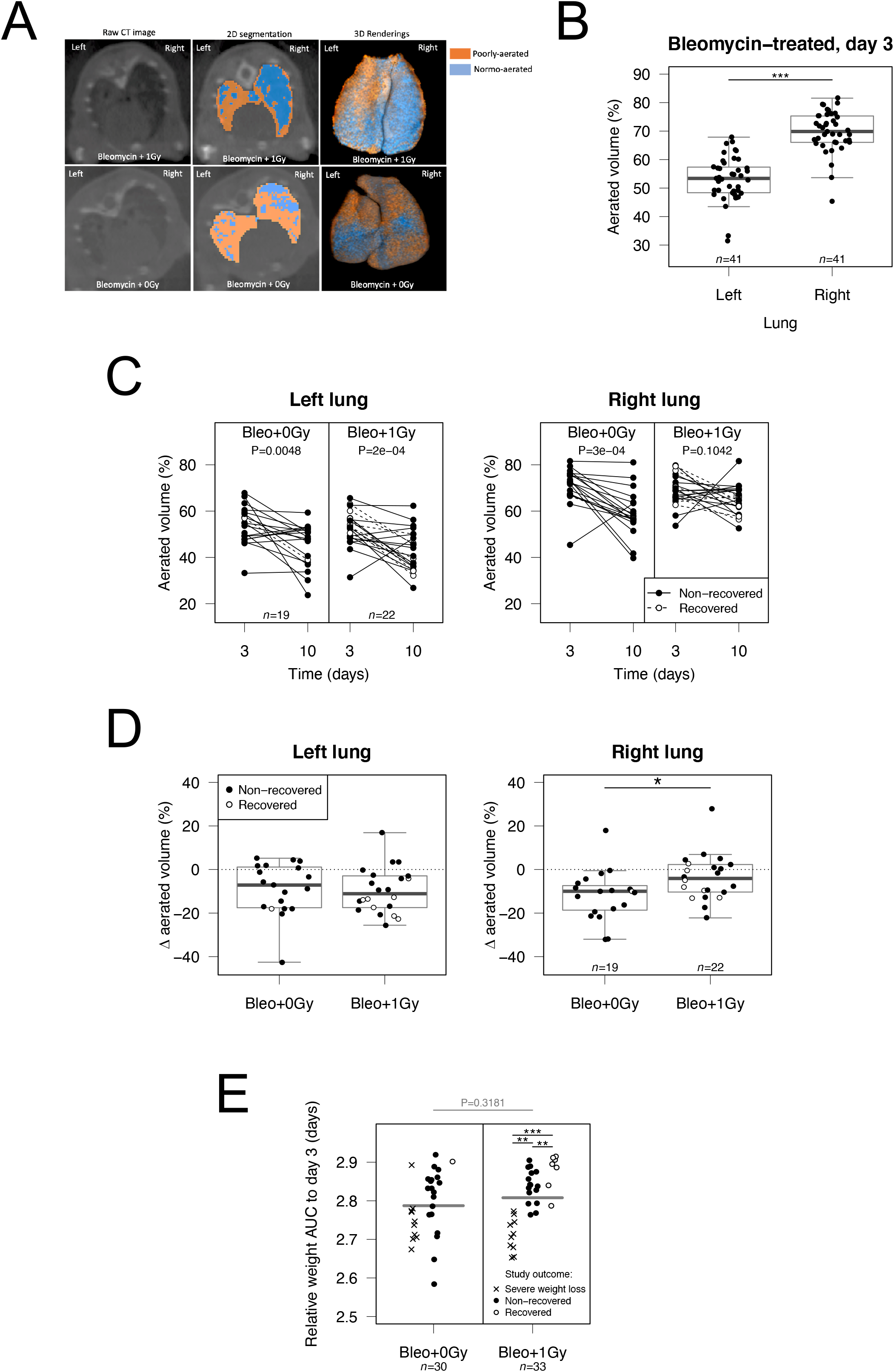
LDLR protects against bleomycin-induced radiological changes in the right mouse lung. (A) The aerated volume of mouse lung was calculated from reconstructed CT images. (B) Aerated volume percentage of each lung in all bleomycin-treated mice at day 3, prior to LDLR. Groups compared by t-test. (C) Percentage aerated volume of each lung on day 3 and day 10. The presence of a consistent trend between time points was assessed by paired t-test. (D) Change in aerated volume percentage between day 10 and day 3 (day 10 – day 3) for each lung. Groups compared by t-test. (E) Relative bodyweight AUC values up to day 3 (prior to LDLR), grouped according to eventual study outcome. Treatment groups compared by t-test and sub-groups compared by one-way ANOVA with *post hoc* Tukey test. Mice whose bodyweight returned to 98% of baseline were classified as recovered. Mice sacrificed early due to an excessive reduction in bodyweight were classified as having experienced severe weight loss. * P<0.05, ** P<0.01, *** P<0.001.

These observations led us to question whether LDLR might be more effective in alleviating moderate (right lungs) than more severe lung pathology (left lungs). To interrogate this further we looked for correlations between the severity of the bleomycin response prior to irradiation (day 3), as indicated by bodyweight AUC, and the likelihood of response to LDLR. Of the mice receiving 1.0 Gy, those that went on to recover had significantly higher AUC values at day 3 than those that did not recover (Fig. 5E). This analysis also confirmed that there was no significant difference in mean pre-LDLR AUC between irradiated and sham irradiated mice, and that mice experiencing a severe initial response to bleomycin (low day 3 AUC) were more likely to go on to experience severe weight loss (humane endpoint), regardless of further treatment.

Finally we evaluated two additional LDLR doses (0.5 and 1.5 Gy) that are also being tested in clinical trials. Neither dose was associated with an improvement in outcome compared to sham irradiation, either in terms of mean bodyweight or likelihood of recovery (Fig. 6A). In keeping with this, no effect of these doses was observed on the immune cell subsets previously shown to respond to 1.0 Gy (Fig. 6B). It should be noted, however, that in this experiment the bleomycin effects were less pronounced than in previous experiments (Fig. 6B, Supp. Fig S8).

**Figure 6:**
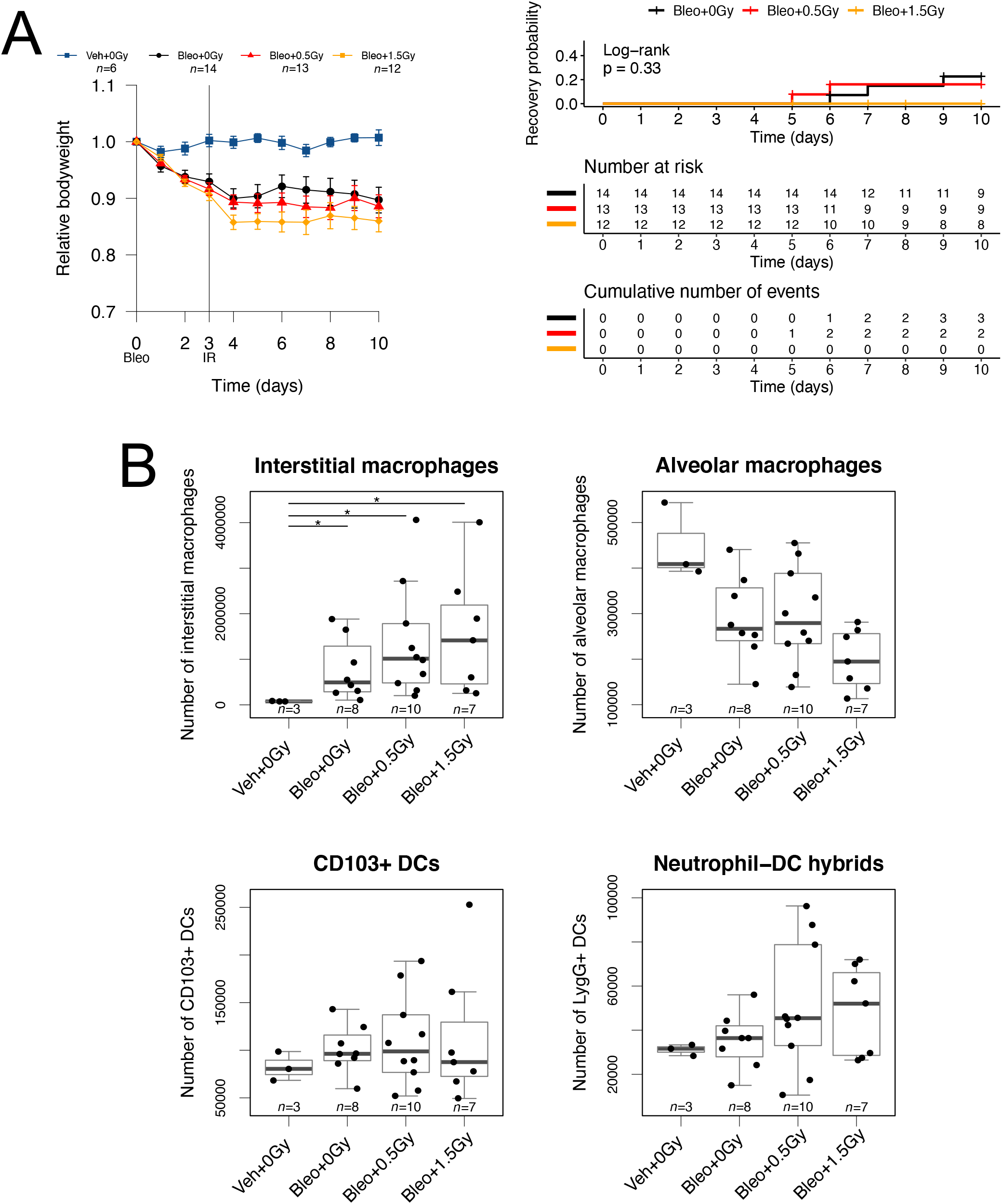
LDLR using 0.5 Gy or 1.5 Gy failed to improve outcomes of bleomycin-treated mice. (A) Relative bodyweight (mean ± SEM) of bleomycin-treated mice following LDLR (0.5 Gy or 1.5 Gy) on day 3. Kaplan-Meier analysis of recovery to 98% of baseline bodyweight. Groups compared by log-rank test. (B) Flow cytometric analysis of immune cells in mouse lung at day 10. Groups compared by Kruskal Wallis and *post hoc* Wilcoxon pairwise test, * P<0.05.

## Discussion

While a substantial body of clinical and preclinical data describes the immunomodulatory effects of low doses of radiation, none of the previously published work has studied pneumonitis. In the absence of a validated small animal model of COVID-19 lung disease (35) and the urgent need for relevant preclinical data, we used the well characterised bleomycin model to undertake pragmatic studies that we hope will provide useful data for clinicians developing early phase studies of LDLR in COVID-19 patients. The bleomycin model was selected because it exhibits many of the pathophysiological changes associated with COVID-19 lung disease, and because recent single cell sequencing studies support the existence of shared immunological mechanisms (21-23). However we recognise its limitations: immune responses to bleomycin and SARS-CoV-2 are not identical, neither within the lungs nor systemically. Furthermore, our experiments were conducted exclusively in female mice aged 11-13 weeks. There is some evidence that young mice are less responsive to bleomycin than older mice (36) and it is possible that male mice would respond differently to bleomycin, LDLR or both. Furthermore, it is well established that the risk of severe COVID-19 lung disease is much greater in older patients (37), and that males are at higher risk of poor outcomes (38).

Having identified bodyweight as a clinically relevant primary endpoint that correlates with the severity of bleomycin-induced pneumonitis and the associated systemic inflammatory response (39), we observed wide variation between mice in terms of rapidity and severity of weight loss, and subsequent recovery. Despite the challenges posed by this variability, our findings support the hypothesis that LDLR, delivered at a time when early histological and immunological features of lung inflammation are apparent, increases the likelihood of recovery in a subset of mice (approximately 25%). Subsequent analyses indicated that mice with moderate lung disease (measured either by lower rates of weight loss or by less marked imaging changes on CT scans) were more likely to respond to LDLR than those with severe pneumonitis. These bodyweight data are supported by histological, radiological and immunological observations which show that LDLR reduces the severity of bleomycin-induced lung changes in a proportion of mice. Of note, histological improvement was observed in a subset of irradiated mice, even though histological assessment was performed exclusively on left lungs, which were generally more severely affected than right lungs. Since the various clinical studies underway are evaluating a range of lung radiation doses from 0.35 to 1.5 Gy, we tested three different doses (0.5, 1.0 and 1.5 Gy) in an attempt to inform clinical decisions in this area. Of these, only 1.0 Gy demonstrated signs of efficacy, although we recognise that the other doses underwent less comprehensive evaluation. This observation is largely consistent with the data generated by Meziani and colleagues in lipopolysaccharide (LPS) and H1N1 influenza models of lung injury, although responses to 0.5 Gy were also observed in some of the histological and cytological readouts reported in this study (11).

Considering possible mechanisms, our cytological studies showed that inhaled bleomycin caused an acute loss of alveolar macrophages and concomitant accumulation of myeloid cell populations in the lung. Similarly, in influenza virus infection, numbers of lung dendritic cells increase as a consequence of more precursor cells migrating to the lung (40). While LDLR was unable to prevent loss of alveolar macrophages, it did reduce accumulation of key dendritic cell, macrophage, and neutrophil populations (Fig. 4). A plausible explanation is that LDLR suppresses the signals that attract precursor dendritic cells and/or inhibits their differentiation into CD103+ dendritic cells. Infection or lung injury can lead to an accumulation of lung macrophages through either recruitment (41) or local proliferation in a Th2-helper environment (42). Bleomycin treatment has been shown to increase production of the chemokine CCL2 by lung cells (43,44); since migration of monocytes into inflamed lungs is dependent on CCL2/CCR2 signalling (43,44) it is reasonable to propose that LDLR might act by reducing CCL2 and/or other signals that attract monocytes into the damaged lung.

Chemokines such as MIP-2 and CXCL5 are released in the first few days after acute lung injury and, together with other factors including extracellular ATP, may play a role in initiating and sustaining accumulation of neutrophils within the lungs following bleomycin inhalation (45). While we saw no increase in classical neutrophils in bleomycin exposed lungs, we did observe accumulation of a hybrid population that expressed markers of both neutrophils (Ly6G) and dendritic cells (CD11c, MHCII). These hybrid cells are thought to differentiate from neutrophil precursors, retaining their phagocytic function while gaining the ability to present antigen to CD4 T cells (33). Inflammation induced by thioglycolate, bacterial or fungal infection leads to an increase in this hybrid population in mouse models of tissue inflammation and these cells have also been found in human tumours (46-48). Our data extend these observations to show that bleomycin also drives accumulation and differentiation of these cells, an effect that we showed to be significantly blunted by LDLR.

### Conclusions

Our data provide preclinical evidence of efficacy of LDLR in a subset of mice with moderate lung injury induced by bleomycin, identify 1.0 Gy as the most effective radiation dose tested, and reveal plausible immunological mechanisms. More comprehensive studies in additional models of pneumonitis and over longer observation periods are warranted to inform ongoing and future studies in COVID-19 patients.

