## Supplementary information for "Low-dose lung radiotherapy for COVID-19 lung disease: a preclinical efficacy study in a bleomycin model of pneumonitis"

**Supplementary data and information.**

### Supp. Figure S1

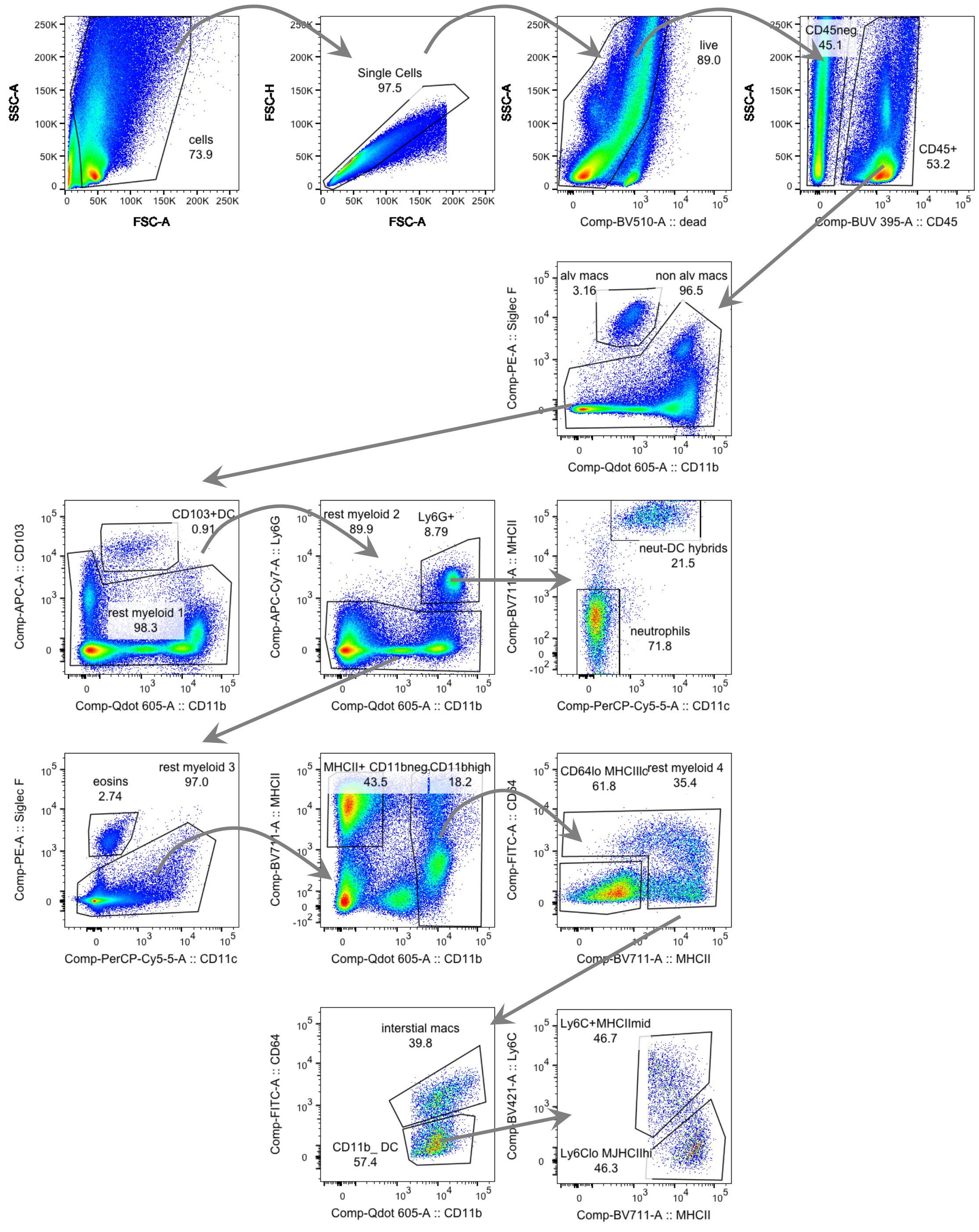


Supp. Figure S1: Gating strategy diagram for flow cytometry analysis of immune cell populations in bleomycin-treated mouse lung.

### Supp. Figure S2

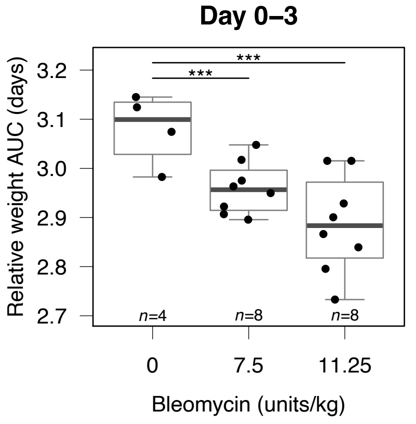


### Supp. Figure S3

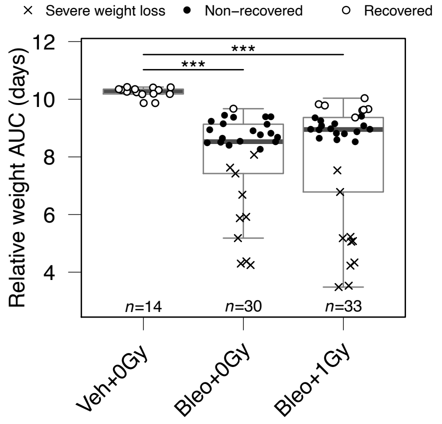


Supp. Figure S2: Relative weight AUC of bleomycin-treated mice to day 3. Groups compared by Kruskal Wallis and *post hoc* Wilcoxon pairwise test, *** P<0.001.

Supp. Figure S3: Relative weight AUC of bleomycin-treated mice to day 10. Mice whose bodyweight returned to 98% of baseline were classified as recovered. Mice sacrificed early due to an excessive reduction in bodyweight were classified as having experienced severe weight loss. Groups compared by Kruskal Wallis and *post hoc* Wilcoxon pairwise test, *** P<0.001.

### Supp. Figure S4

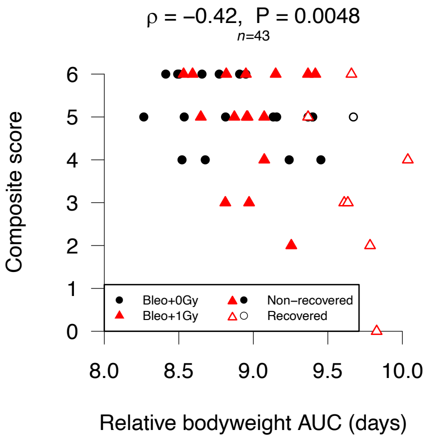


### Supp. Figure S5

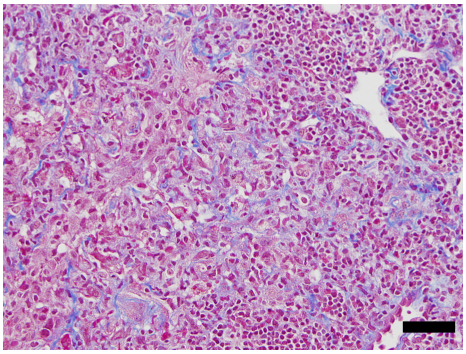

Supp. Figure S4: Spearman correlation of immune infiltrate histological composite score and relative bodyweight AUC. Mice whose bodyweight returned to 98% of baseline were classified as recovered.

Supp. Figure S5: Pulmonary histology of a sham-irradiated mouse 10 days after bleomycin-treatment. Fine wispy blue material depicts early aberrant collagen deposition (Masson’s Trichrome stain. Scale bar: 50 μm ).

### Supp. Figure S6

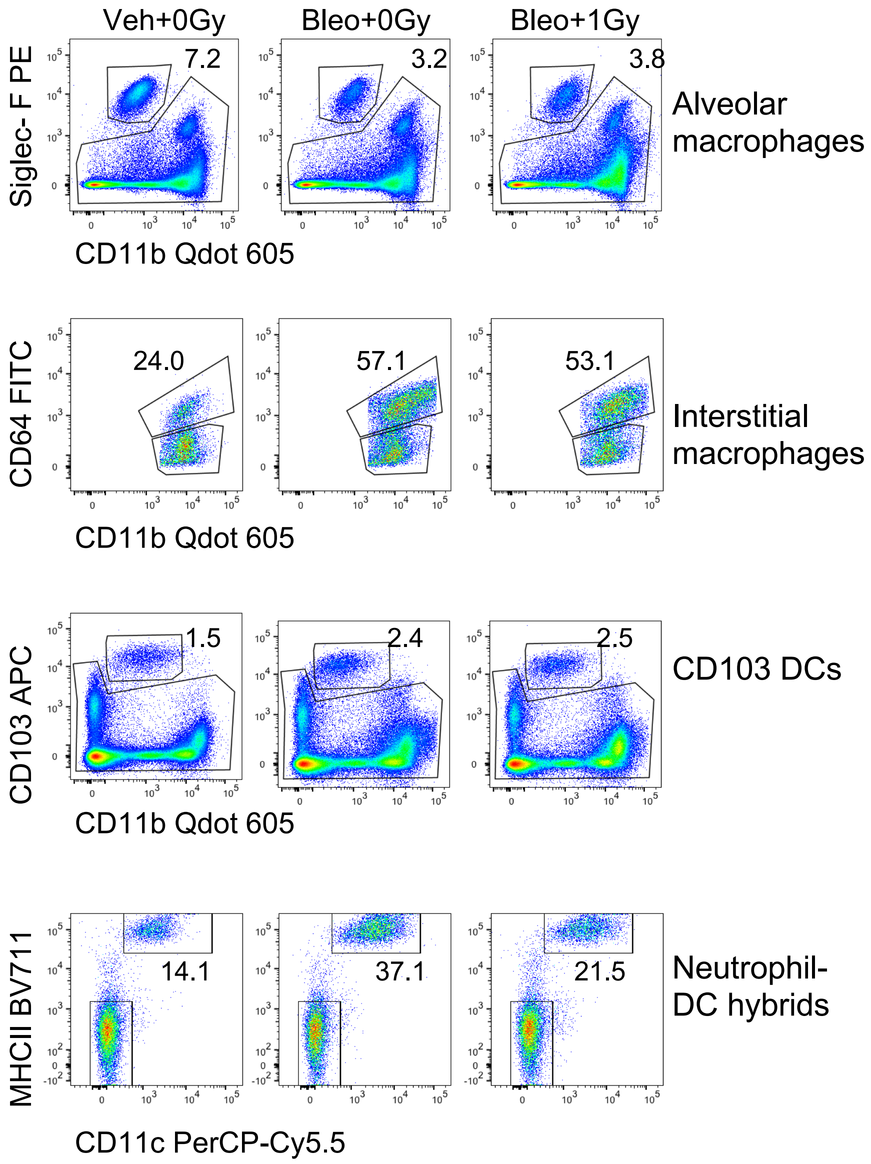


Supp. Figure S6: Example flow cytometry plots for the quantification of immune cell populations from bleomycin-treated mouse lung.

### Supp. Figure S7

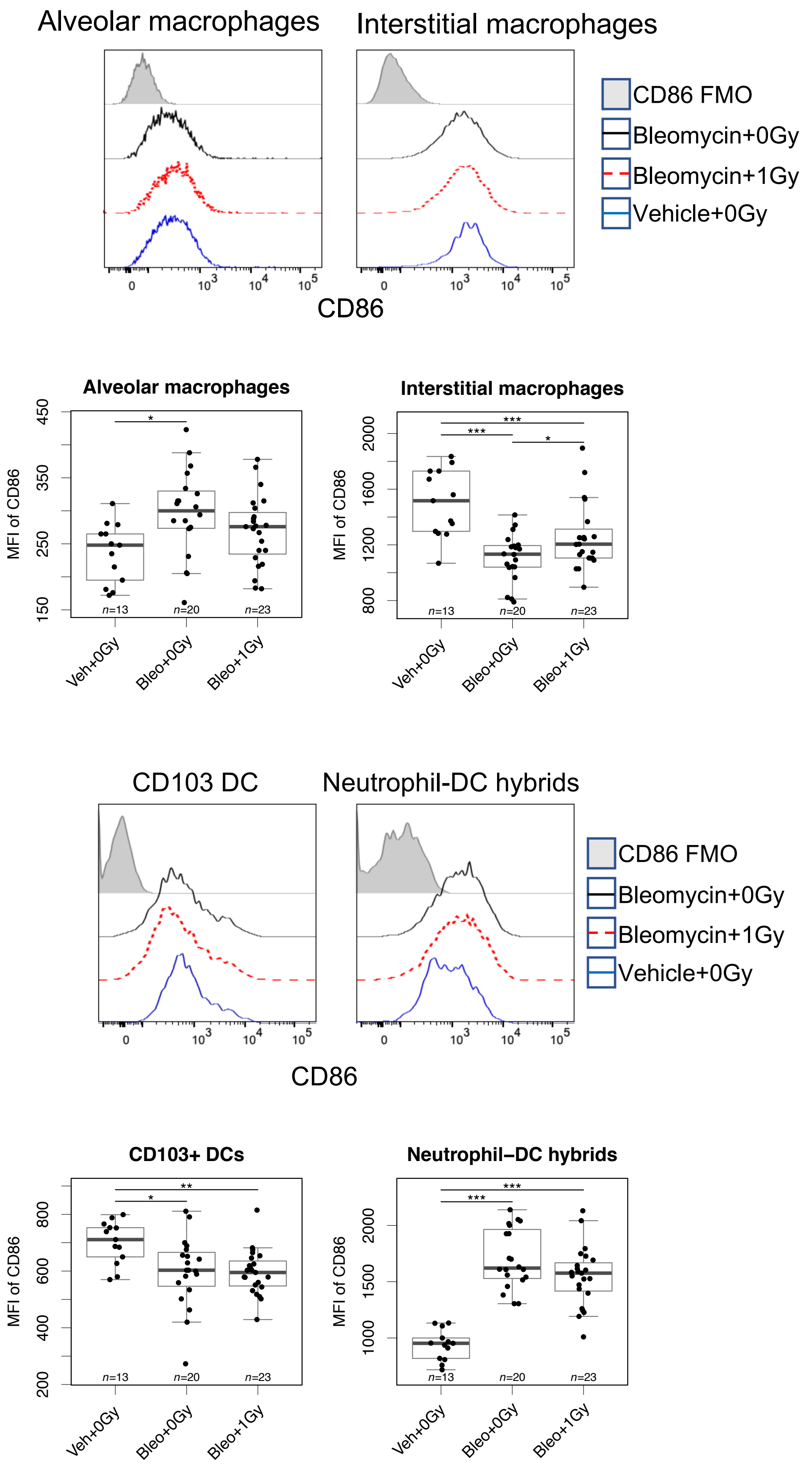


Supp. Figure S7: CD86 expression by immune cell populations of interest. FMO – fluorescence minus one control, MFI – mean fluorescence intensity. Groups compared by Kruskal Wallis and *post hoc* Wilcoxon pairwise test, * P<0.05, ** P<0.01, *** P<0.001.

### Supp. Figure S8

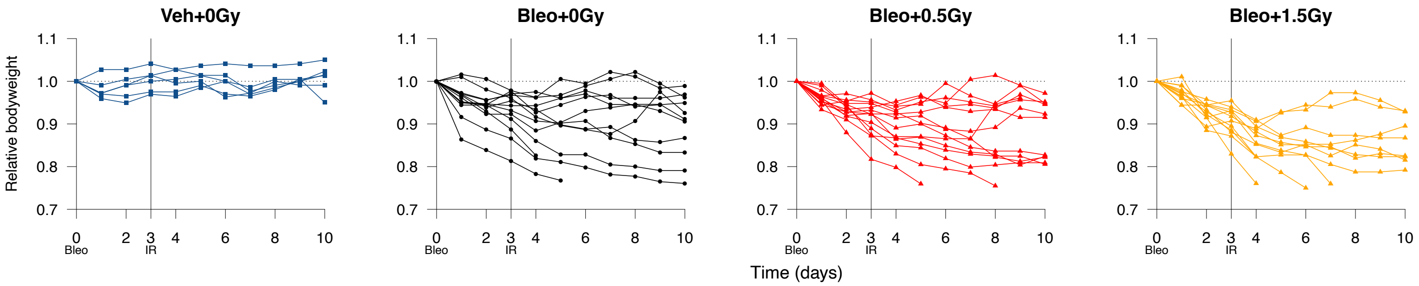


Supp. Figure S8: Relative bodyweight of individual mice treated with bleomycin and LDLR at 0.5 Gy or 1.5 Gy.

### Supp. table S1

Supp. Table S1: Antibodies used for flow cytometry.

| **Fluorochrome** | **Ab** | **Company** | **Clone name** |
|---|---|---|---|
| FITC | CD64 | BioLegend | X54-5/7.1 |
| PerCP-Cy5.5 | CD11c | BioLegend | N418 |
| APC-Fire750 | Ly6G | BioLegend | 1A8 |
| APC | CD103 | BioLegend | 2E7 |
| BUV395 | CD45 | BD | 30-F11 |
| BV785 | B220 | BioLegend | RA3-6B2 |
| BV711 | MHCII | BioLegend | M5/114.15.2 |
| BV605 | CD11b | BioLegend | M1/70 |
| BV421 | Ly6C | BioLegend | HK1.4 |
| PeCy7 | CD86 | BioLegend | GL-1 |
| PE | SiglecF | BioLegend | S17007L |

#### Supplementary information

##### Area-under-the-curve (AUC)

Area-under-the-curve (AUC, linear interpolation) analysis of mouse daily bodyweight was used to quantify the response to bleomycin over time, in a single parameter. Since AUC was less influenced by day-to-day mouse bodyweight variation than individual daily weights, it was used firstly to exclude mice that had been largely unaffected by bleomycin administration. As depicted in Figure 1E, mice were classified as a bleomycin responder or non-responder based on their day 3 AUCs. A standardised day 3 AUC threshold for inclusion/exclusion of 2.92 was derived from the responses of naïve and vehicle-treated mice in pilot studies and used for all subsequent mouse cohorts.

Secondly, analysis of bodyweight AUC across the entire time-course of the experiment was performed. This approach had the advantage of representing all mice, including those that exhibited severe weight loss and hence were humanely culled before the end of the study. Through comparison with vehicle-treated control animals, mice showing recovery of bodyweight were readily identifiable. In turn, representation of the bodyweight response over time as a single value allowed further analyses to be performed, such as correlation with other biological measures (for example Supp. Figure S4).

##### Recovery probability

As reported, LDLR showed efficacy in only a proportion of bleomycin-treated mice. As a result, the existence of an LDLR-responsive subset of mice may be masked by commonly-used statistical analyses, for example when comparing group means (Figure 2A). Accordingly, for our study we employed Kaplan-Meier analysis, which is more suited to this scenario (hence its use for clinical datasets), to interrogate the efficacy of LDLR.

For this, we recorded the time taken for the bodyweight of individual mice to exceed a recovery threshold for at least one day after receiving LDLR. Mice that were culled early due to severe weight loss were censored at the appropriate time in the analysis, as they were no longer ‘at risk’ of recovery. Thus, both the proportion and the timing of recovery events informed the P-value generated by log-rank testing, under the null hypothesis that there was no difference between the populations in terms of their probability of recovery.

In this type of analysis, the outcome is naturally heavily influenced by the definition of recovery used. To increase transparency, we present results determined using three different recovery thresholds. Intuitively, recovery to 100% of baseline bodyweight was selected, alongside recovery to 96% based on the behaviour of vehicle-treated control mice (see Figure 2B). An intermediate definition (recovery to 98% of baseline bodyweight) was presented as our primary readout.

##### Scoring lung tissue inflammation

Since bleomycin was administered intranasally, the earliest and most substantial changes were typically observed close to the larger airways. Therefore, H&E-stained

sections were screened for suitability before scoring and semi-quantitative scores were based on the most substantially affected lung regions to reflect this intentional sampling bias. This has been accepted as the best approach for scoring for the inherent patchy distribution of lesions observed in the bleomycin model (1).

The macrophage 5-point scoring system, based on scoring protocols for fibrosis following administration of bleomycin (2), was used with a minimum score ‘0’ corresponding to no significant changes and a maximum score ‘4’ representing full obliteration of lung tissue by macrophage infiltrates. An intermediate score ‘2’ reflected developing macrophage infiltrates between the extremes, whilst nuances were scored as ‘1’ or ‘3’.

Perivascular/peribronch(iol)ar lymphocytes forming well-appreciable aggregates (observed at 4x low magnification) were scored as ‘1’ to represent rare lymphoid nodules or small numbers of lymphocytes in multiple nodules, as ‘2’ to represent larger numbers/aggregates of lymphocytes, or as ‘0’ to represent an absence of lymphocyte aggregates at this magnification.

Macrophage and lymphocyte scores were summed to provide a composite score.

##### Lung CT quantification

Previous studies have shown the value of using CT for longitudinal evaluation of lung disease in the mouse bleomycin model. Since bleomycin is administered via the airways, differences in lesion distribution between lungs may be observed, as in our study, and this can be appreciated and quantified on CT imaging. A semi-automatic lung segmentation method was used to define whole lung volumes; the lung parenchyma was easily detectable and well defined by HU thresholding. All segmentations were visually inspected and manually corrected if considered necessary. Poorly aerated lung volume was used as a marker of lung disease as previously reported (3).


1. Jenkins RG, Moore BB, Chambers RC, Eickelberg O, Konigshoff M, Kolb M*, et al.* An Official American Thoracic Society Workshop Report: Use of Animal Models for the Preclinical Assessment of Potential Therapies for Pulmonary Fibrosis. Am J Respir Cell Mol Biol **2017**;56:667-79
2. Hubner RH, Gitter W, El Mokhtari NE, Mathiak M, Both M, Bolte H*, et al.* Standardized quantification of pulmonary fibrosis in histological samples. Biotechniques **2008**;44:507-11, 14-7
3. Ruscitti F, Ravanetti F, Essers J, Ridwan Y, Belenkov S, Vos W*, et al.* Longitudinal assessment of bleomycin-induced lung fibrosis by Micro-CT correlates with histological evaluation in mice. Multidiscip Respir Med **2017**;12:8
